# Soil nutrient stoichiometry affects the initial response of microbial community to trophic perturbation

**DOI:** 10.1101/323709

**Authors:** Kazumori Mise, Runa Maruyama, Yuichi Miyabara, Takashi Kunito, Keishi Senoo, Shigeto Otsuka

## Abstract

Soil microbes are drivers of global ecosystem functionality and are continuously subjected to external perturbations. It is fundamental for ecologists and environmental scientists to understand and further predict the microbes’ responses to these perturbations. A major and ubiquitous perturbation is the addition of chemical nutrients, including fertilizers and animal urine, to soil. Recent biogeographical studies suggest that soil nutrient stoichiometry (i.e., nutritional balance) determines microbial community structure and its functions with regard to material circulation. Given this information, here, we show that soil nutrient stoichiometry, or the bioavailable C:P ratio, determines the impact of nutrient addition on the soil’s microbial communities. We sampled two soils with similar carbon and nitrogen concentrations but with a 20-fold difference in phosphorus bioavailability. Soil microcosms with carbon and nitrogen amendments were constructed for both the soils. The phosphorus-depleted soil received prolonged effect from carbon and nitrogen amendments: the phosphatase activity gradually increased over a 24-day incubation period and the microbial community structure did not present recovery to its initial state. In contrast, in the other soil, both phosphatase activity and microbial community structure gradually returned to those of the control samples. Phosphorus depletion mitigated carbon and nitrogen intake; therefore, the effects of carbon and nitrogen amendment lasted longer. Our results demonstrate that nutritional stoichiometry is a strong predictor of microbial community dynamics in response to trophic perturbation, particularly when considering the length of time the trait of perturbation persists in the soil.

## 1. Introduction

Soil microbial communities play the crucial roles in many ecological functions. They are the drivers of soil nutrient cycling, primary production, and decomposition of macromolecules, all of which are essential for the integrity of global ecosystem. Thus, the maintenance of community functionality is of great interest to us.

Soil microbial communities are subject to a wide variety of external perturbations, and adaptively respond to the environmental changes initiated by the perturbation (Griffith et al., 2013). They undergo drastic changes both in composition and function immediately after perturbation (Knelman et al., 2014; Castle et al., 2017), and primary succession determines the subsequent trajectory of microbial community assembly (Pagaling et al., 2014). Hence, formulating the early responses of soil microbial community structure and function to perturbation is a fundamental focus in soil microbial ecology.

One prominent perturbation that soil receives is the input of soil nutrition such as bioavailable carbon (C) and nitrogen (N) sources. The sources of nutrient input are highly diverse, including anthropogenic fertilization practices on arable soils, litter supply on forest soils (Wang et al., 2014), and mammalian urine deposition on grassland soils (Buscardo et al., 2017). Such nutritional input has been demonstrated to alter soil microbial community structure and function; for example, microcosm experiments have shown that nutrition input results in higher microbial respiratory activity (Castle et al., 2017). Additionally, microbes with high copy number of rRNA gene are shown to dominate soils immediately after soil nutritional inputs (Nemergut et al., 2016). However, no uniform discipline has been established to date regarding the community dynamics in response to nutrient input.

Soil nutrient status is often described by the idea of nutrient stoichiometry. For example, soil C:N ratio has commonly been used as a criterion of soil fertility and the soil microbial activity, in the context of agricultural studies. Besides, soil nutrient stoichiometry has been shown to drive microbial communities both in terms of structure and function. One recent study suggests that the soil C:N:P ratio is strongly correlated with the bacterial community structure in soil, presumably because high C:P and N:P ratios induce competition among bacteria, which may result in the extinction of bacteria intolerant of phosphorus (P) scarcity (Delgado-Baquerizo et al., 2017). Regarding microbial community function, the resource allocation model on the microbial production of soil extracellular enzymes (Sinsabaugh and Moorhead, 1994) is noteworthy. It claims that soil microbes optimize C-, N-, and P-acquiring enzyme syntheses to maximize their growth, in accordance with soil nutrient stoichiometry. This model is empirically supported by Fujita et al. (2017), who reported that the ratio of P- and C-acquiring enzyme activities were negatively correlated with the concentration of soil bioavailable phosphorus.

Although these insights suggest that nutrient stoichiometry drives the soil microbial community structures and functions, all of the above-mentioned studies deal with communities in its stable states. Thus it remains unknown whether nutrient stoichiometry is a strong predictor of soil microbial community dynamics in response to trophic perturbation (i.e. nutrient input), which we aimed to testify in the present study. As a model to address this question, we focused on P-limited conditions induced by C and N input. P-limitation is often observed in terrestrial ecosystems, and abovementioned studies suggest that soil P content or P availability drives microbial community structure and functionality (Delgado-Baquerizo et al., 2017; Fujita et al., 2017). We hypothesized that stronger phosphorus limitation (i.e. more unbalanced C:P stoichiometry) results in more drastic changes in microbial community structure and functionalities. To test this hypothesis, we performed soil incubation experiments using two soils differing in available phosphorus concentration but with similar CN content. We loaded labile C and N sources to soil microcosms, and monitored the soil nutritional stoichiometry, potential phosphatase activity and prokaryotic community structure for 24 days.

## 2. Materials and methods

### 2.1 Construction of soil microcosms

Soil samples used in this study were arable Andisols (hereafter referred to as “soil X”) and brown forest soils (“soil Y”) sampled in Nagano Prefecture, Japan. Detailed sampling sites and chemical properties are indicated in Table 1. Soil C:N ratios were mostly the same within the two soils, while the C:P ratio of soil Y was 5 or 10 times smaller than that of soil X.

**Table 1.** Sampling locations and chemical properties of the two soils.

|  | Soil X | Soil Y |
| --- | --- | --- |
| Sampling location | 36.10°N, 137.93°E | 36.28°N, 137.99°E |
| Soil type | Andisol | Brown forest soil |
| Land use | Arable soil | Forest |
| pH(H<sub>2</sub>O) | 6.90 | 5.38 |
| DOC (mg kg<sup>-1</sup>-drysoil) | 230 | 207 |
| Exchangeable Mineral N (mg kg<sup>-1</sup>-drysoil) | 38.4 | 16.0 |
| Total C (g kg<sup>-1</sup>-drysoil) | 39.5 | 30.0 |
| Total N (g kg<sup>-1</sup>-drysoil) | 2.9 | 2.1 |
| Truog-P (mg kg<sup>-1</sup>-drysoil) | 244 | 20.6 |
| H<sub>2</sub>O-P (mg kg<sup>-1</sup>-drysoil) | 22.3 | 5.06 |
| Total C / Total N | 13.6 | 14.3 |
| Truog-P / Total C (×10<sup>-4</sup>) | 61.7 | 6.87 |
| H<sub>2</sub>O-P / DOC (×10<sup>-4</sup>) | 5.65 | 1.69 |

For each soil, 93 microcosms, consisting of approximately 15 or 30g of soil on Petri dishes, were constructed. After five days of pre-incubation at 23 °C without light, three microcosms were destructively sampled. To 72 samples out of the remaining 90, a carbon (C) source (glucose or cellobiose solution) and nitrogen (N) source (ammonium chloride solution) were amended. Half of the microcosms were amended with a quarter amount of C and N at the beginning of incubation, and received additional amendment every seven days. The total amount of amended C and N were same in all samples after 24 days of incubation. Besides the negative controls (no CN amendment), four types of treatment in total were prepared, representing two types of C source and two patterns of CN amendment: glucose + NH_4_Cl amended at one time (hereafter abbreviated as G-1); cellobiose + NH4Cl amended at one time (C-1); glucose + NH_4_Cl separately amended (G-4) and cellobiose + NH_4_Cl separately amended (C-4) (Table 2).

**Table 2.** Details in the carbon and nitrogen amendment to soil microcosms. “Day 0” is defined as the time incubation was started.

| Treatment | Amended carbon and nitrogen sources<br>(per 1g of wetsoil) | Timing(s) of amendment |
| --- | --- | --- |
| Control | – | – |
| G-1 | Glucose (8.4 mg-C) + NH<sub>4</sub>Cl (420 µg-N) | Day 0 |
| C-1 | Glucose (8.4 mg-C) + NH<sub>4</sub>Cl (420 µg-N) | Day 0 |
| G-4 | Cellobiose (2.1 mg-C) + NH<sub>4</sub>Cl (105 µg-N) | Days 0, 7, 14 and 21 |
| C-4 | Cellobiose (2.1 mg-C) + NH<sub>4</sub>Cl (105 µg-N) | Days 0, 7, 14 and 21 |

The 90 microcosms were incubated at 23 °C and destructively sampled after 24 hours and 3, 10, 17, 23, and 24 days of incubation. Three microcosms per each treatment type were sampled on each of these days. Here we designed destructive sampling experiments to make each microcosm small and thereby mitigate the experimental deviation that results from heterogeneity within one microcosm.

### 2.2 Soil chemical/biochemical analyses

The concentrations of total C and N (TC and TN) were measured using a Nitrogen and Carbon analyzer (Thermo Finnigan Flash EA1112; Thermo Fisher Scientific, Waltghram, MA, USA). Soil samples with 0, 3, 10, 17 and 24 days of incubation were subjected to enzyme activity measurement and prokaryotic community analysis. Alkaline phosphatase (ALP), acid phosphatase (ACP), and β-D-glucosidase (BG) activities were determined as described in Tabatabai (1994). Briefly, ALP and ACP activities were measured in modified universal buffer (MUB) at pH values of 11 and 6.5, respectively, using *p*-nitrophenyl phosphate as a substrate. The β-D-glucosidase activity was measured in MUB at pH 6.0, using *p*-nitrophenyl-β-D-glucopyranoside as a substrate. Additionally, pH of soil samples incubated for 0, 3 and 24 days were measured using a glass electrode in 1:2.5 (w/v) soil-water suspension.

Soil samples with 24 hours and 23 days of incubation were subjected to the quantifications of dissolved organic carbon (DOC), inorganic nitrogen and bioavailable phosphorus. DOC was extracted in 0.5 M K_2_SO_4_ solution, and the concentration was determined using a TOC meter (TOC-5000; SHIMADZU, Kyoto, Japan). Bioavailable phosphorus was independently extracted and quantified in two ways: extraction in H_2_SO_4_ solution (Truog, 1930; Truog–P) and water extraction (H_2_O–P). Concentrations of inorganic P (Pi) in the extracts were determined using the ammonium molybdate-ascorbic acid method (Murphy and Riley 1962).

### 2.3 Amplicon sequencing of the 16S rRNA gene

Soil DNA was extracted from 380–420 mg of soil using FastDNA SPIN Kit for Soil (Qbiogene, Carlsbad, CA, USA) for soil X and ISOIL for beads beating (Nippon Gene, Toyama, Japan) for soil Y. To improve the yield of DNA from soil X, autoclaved 2% (w/v) casein solution (dissolved in 300mM sodium phosphate, pH 8.0) was added. The amplicon sequencing of the 16S rRNA gene was performed as described by Caporaso et al. (2012). Briefly, the V4 region of the 16S rRNA gene was amplified using 515F/806R primers. The PCR products were subjected to Illumina sequence-by-synthesis on MiSeq Reagent Kit v2 (Illumina, San Diego, CA, USA).

The nucleotide sequences obtained were quality-filtered with USEARCH ver 9.2.64 (Edgar 2010, Bioinformatics) and subjected to operational taxonomic unit (OTU) clustering, at a similarity threshold of 97% or more, with UPARSE (Edgar, 2013 Nat Methods). Each OTU was taxonomically annotated using the RDP Classifier (Wang et al., 2007 AEM) and Greengenes 13_8 (McDonald et al., 2012) implemented in MacQIIME ver 1.9.1 (Caporaso et al., 2010b), with a confidence threshold of 0.8. OTUs annotated as organellar ribosomal genes or unclassified at the domain level were discarded. A phylogenic tree of the OTU representative sequences was constructed using PyNAST (Caporaso et al., 2010a) and FastTree (Price et al., 2009). The number of sequences was normalized to 15,946 reads per each sample by random sampling. Subsequently, Simpson’s diversity index of each community and weighted UniFrac distances (Lozupone and Knight, 2005) between communities were calculated with MacQIIME ver 1.9.1.

Additionally, average rRNA gene copy number was calculated as described by Nemergut et al. (2016). Briefly, the OTU representative sequences were mapped to rRNA gene copy number database packaged in PICRUSt (Langille et al., 2013).

### 2.4 Statistical analyses

Two-dimensional non-metric dimensional scaling (2D-NMDS) was performed based on UniFrac distances using MacQIIME ver 1.9.1. To explore OTUs that showed characteristic transition patterns in microbial community, SIMPER analysis was performed. Additionally, ANOVA was employed to assess the significance of CN treatment in determining soil chemical characteristics and enzymatic activities. R 3.4.0 (R core team, 2017) was used for all the statistical analysis except for 2D-NMDS.

## 3. Results

### 3.1 Soil nutrient and enzymatic stoichiometry

As a whole, the microcosm groups of two different soil types presented contrasting trends regarding nutrient availability. Changes in soil chemical properties varied between soil types, amendment patterns, and the types of amended carbon sources. After 24 hours of incubation, DOC concentrations in soil X were 279–2270mg kg^-1^-drysoil (recovery rate: 7.30–26.2%), significantly lower than those in soil Y (739–5900mg kg^-1^-wetsoil, recovery rate: 32.4–68.8%) (Fig. 1a, 1c). The DOC concentrations declined after 23 days of incubation (Fig. 1b, 1d).

**Figure 1.**
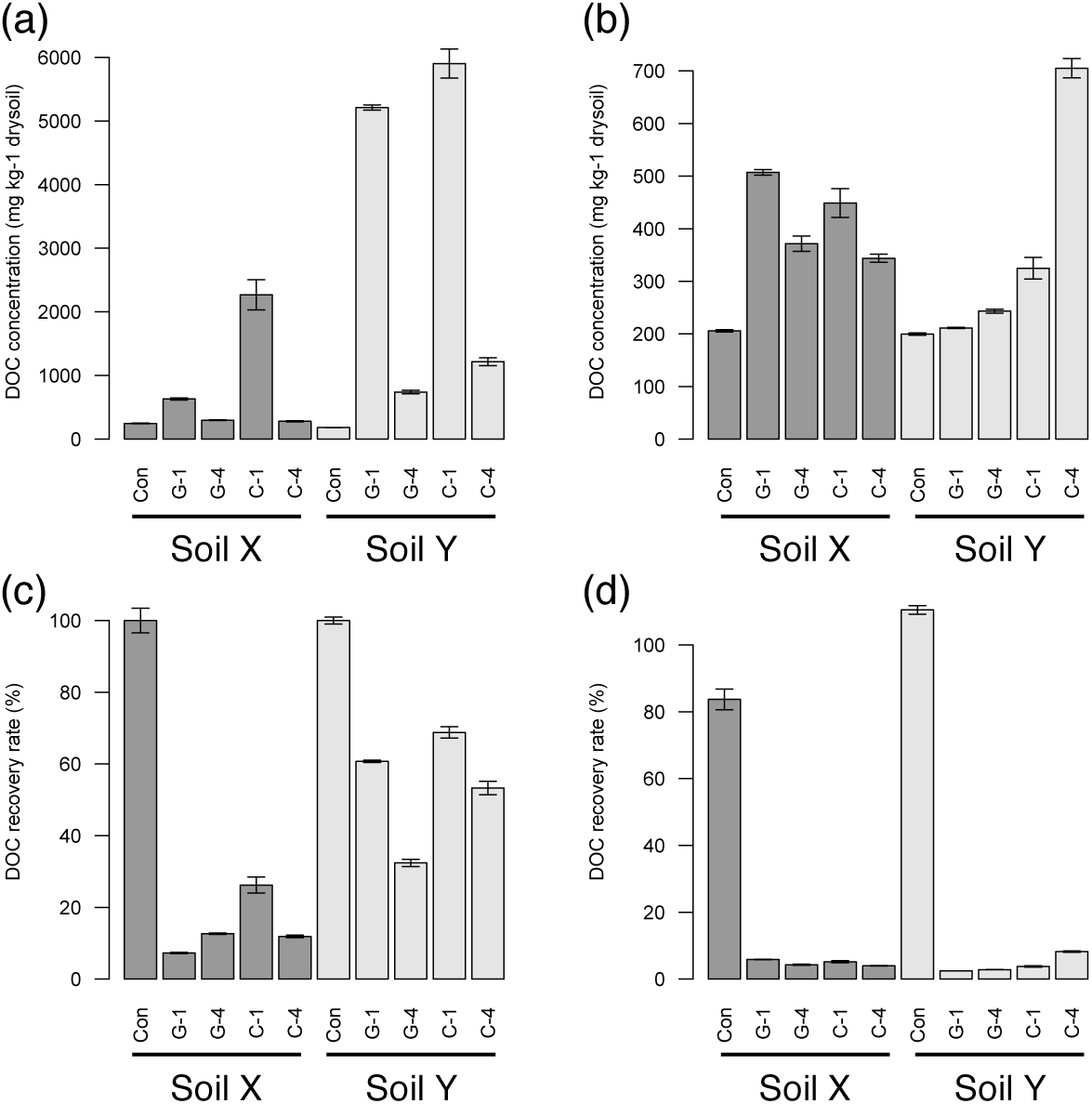
DOC concentration (a,b) and DOC recovery rate (c,d) in soil microcosms after 24 hours of incubation (a,c) and 23 days of incubation (b,d). DOC recovery rate is calculated as follows: (DOC recovery rate) = (DOC concentration)/(initial DOC concentration + amount of amended carbon). We regarded initial DOC concentration equal to the DOC concentration of control samples after 24 hours of incubation. “Con” stands for “Control”.

### 3.2 Soil enzyme activities

Soil ALP and ACP activities were precipitously elevated after the initial carbon and nitrogen amendment (Fig. 2). Overall, ALP had higher activity than ACP in soil X, and vice versa in soil Y. While intensely-disturbed groups of soil X presented decreases in ALP and ACP activity, those of soil Y maintained the phosphatase activity even after 24 days of incubation. Notably, phosphatase activity in soil Y was marginally affected by the amount of carbon input, compared with soil X (Fig. 3).

**Figure 2.**
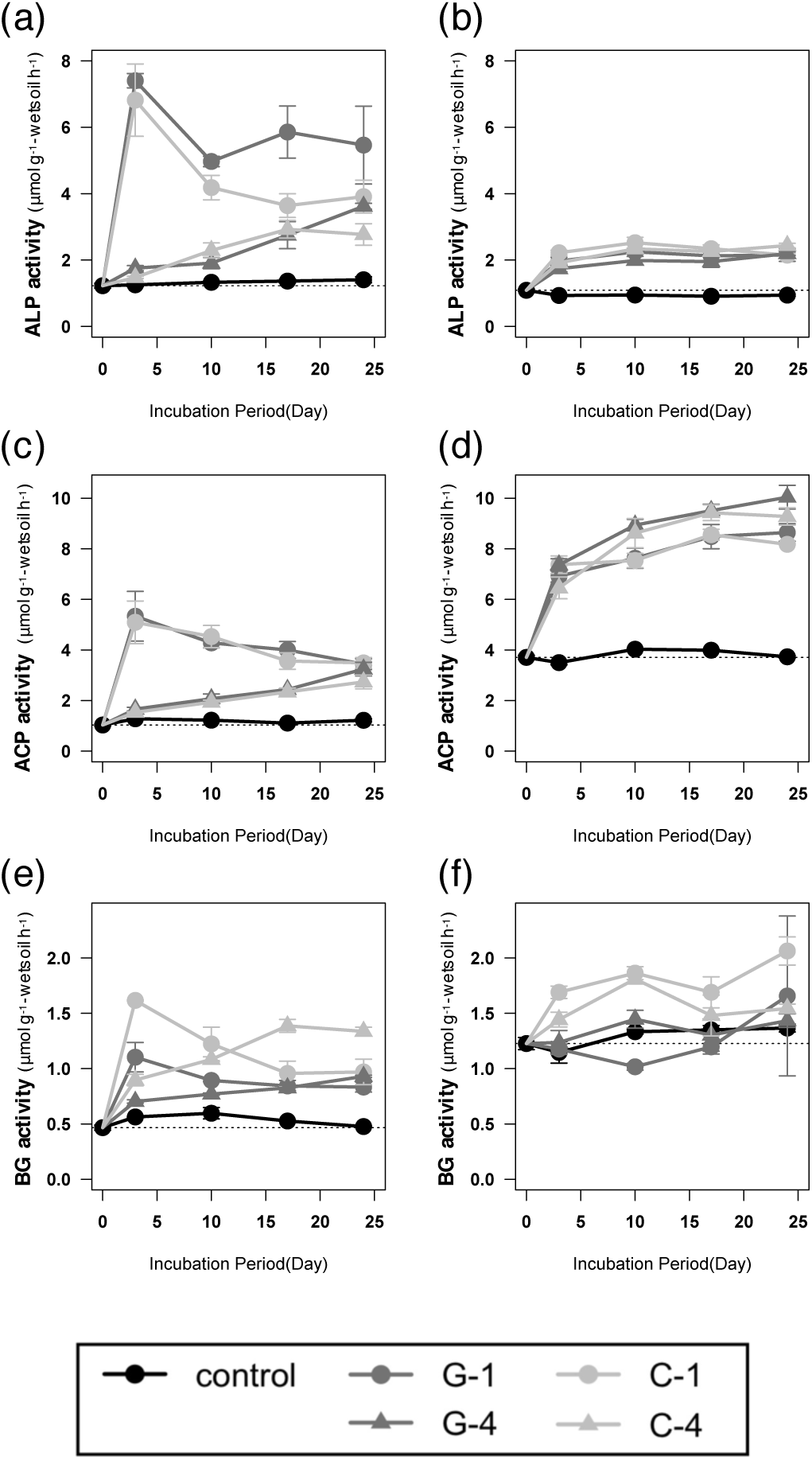
ALP (a,b), ACP (c,d) and BG (e,f) activities of soil X (a,c,e) and soil Y (b,d,f) in the course of 24 days of incubation.

**Figure 3.**
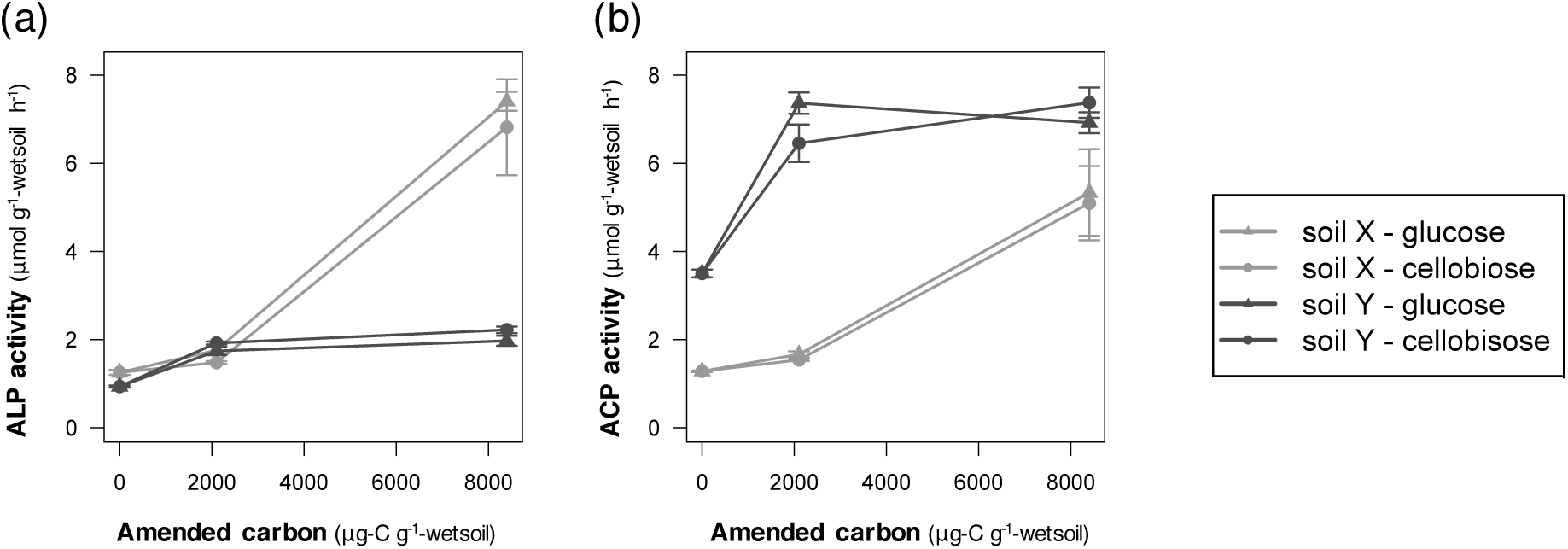
Relationships between ALP (a) and ACP (b) activities at day 3 and amount of amended carbon at the beginning of incubation.

With respect to the kind of carbon source amended, higher BG activity was observed in the cellobiose-amended group than in the glucose-amended group. Conversely, the relationship between phosphatase activity and carbon amendment was not consistent. ALP activity in soil X and ACP in soil Y were significantly higher under glucose amendment, while ACP in soil X and ALP in soil Y were not.

### 3.3 Prokaryotic community structure

A total of 2,514,047 high-quality sequences were obtained and 7,968 non-chimeric OTUs were defined. The rarefaction curves suggest that most prokaryotic classes were detected, although some rarebiosphere OTUs were left undetected (Fig. S1).

Microbial community evenness was estimated using the Simpson index. In soil X, Simpson indices sharply increased after the initial CN input, and gradually dropped over the course of 24 days (Fig. 6a). By contrast, in the G-1 and C-1 groups of soil Y, the Simpson indices continuously increased until 17 days after the initial CN input (Fig. 6b).

**Figure 4.**
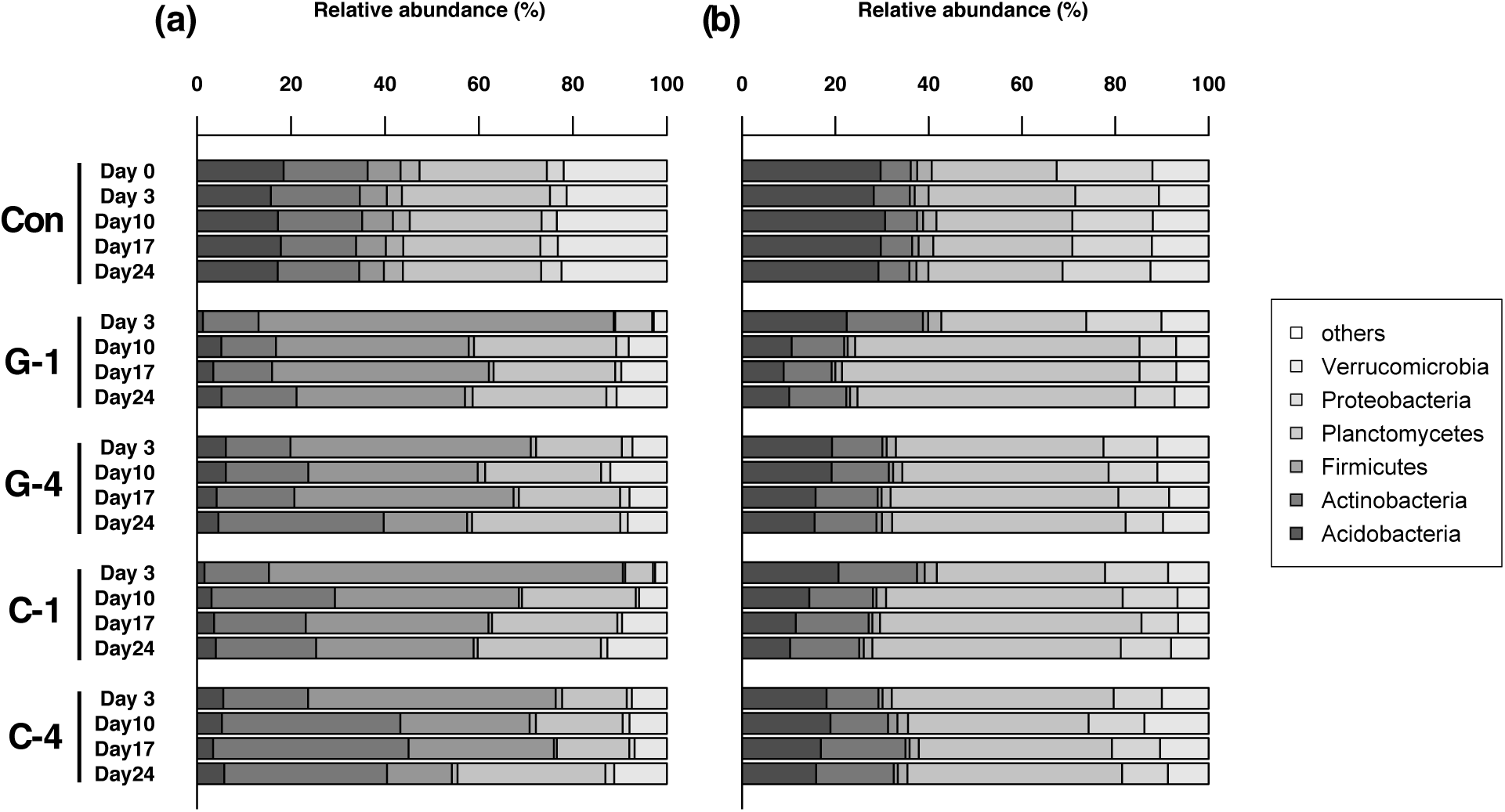
Prokaryotic community structure of incubated soil X (a) and soil Y (b). Presented are the average of three replicates. “Con” stands for “Control”.

**Figure 5.**
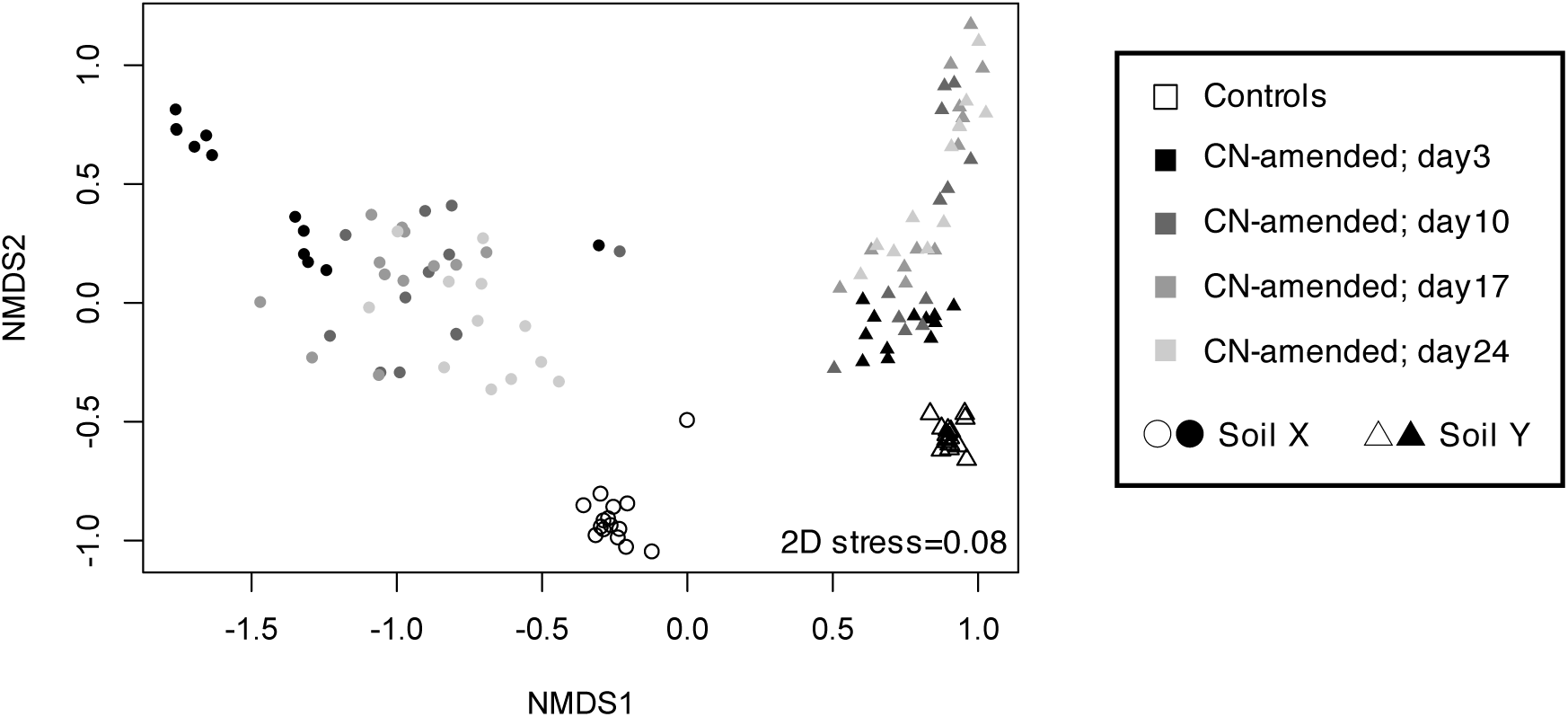
Transition in microbial community structure during the 24 days of incubation, illustrated by 2D-NMDS based on weighted UniFrac distances.

**Figure 6.**
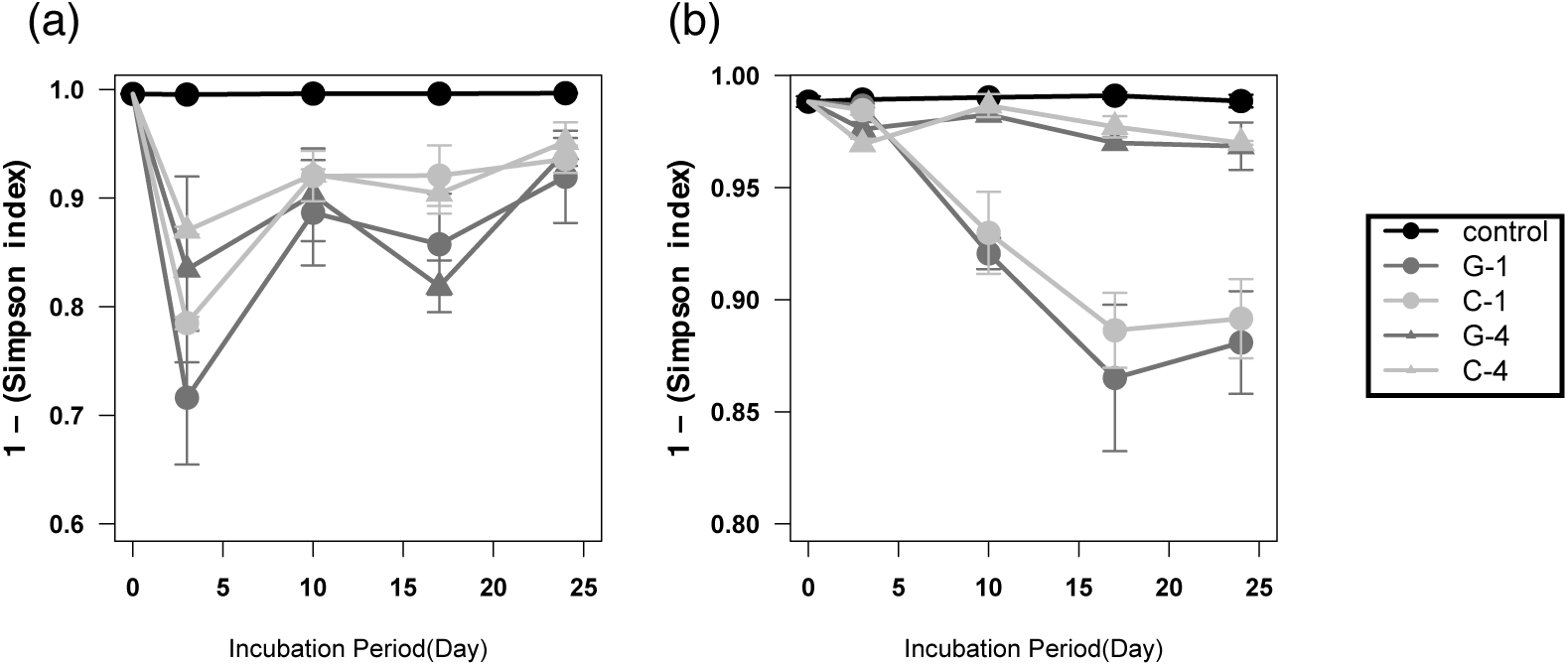
Transition of Simpson index (complementary) in soil X (a) and soil Y (b). Presented are the average of three replicates.

The proportion of the phyla *Firmicutes* and *Proteobacteria* was precipitously elevated after the initial perturbation in soil X and Y, respectively (Fig. 4). Subsequent transitions were subtler than in the first three days, although a gradual change was observed in all but the control group. Over the course of 24 days of incubation, the proportion of the phylum *Firmicutes* in soil X decreased, while the proportion of the phylum *Proteobacteria* in soil Y did not. In line with this observation, the entire compositions of the microbial communities returned to the initial state (or those of the control samples) in soil X, while they did not in soil Y (Fig. 5). Via dissection at individual OTU levels using SIMPER analysis, we identified OTUs that were specifically affected by each treatment: OTUs annotated as the genus *Bacillus* and the genus *Arthrobacter* (phylum *Actinobacteria*) in soil X, and one annotated as the genus *Rhodanobacter* (class *Gammaproteobacteria*) in soil Y, were found to be prominent drivers of community transitions (Fig. S2a). Notably, OTU_3 (genus *Rhodanobacter*) in soil Y gradually increased throughout the course of 24 days of incubation, while OTU_1 (genus *Bacillus*) and OTU_2 (genus *Arthrobacter*) in soil X did not (Fig. S2b–d).

The average rRNA gene copy number exhibited sharp growth during the first three days of incubation in both soil X and soil Y (Fig. 7). After three days of incubation, the average copy number was up to 7 copies/cell in soil X, more than three times larger than those in soil Y. Subsequently, the average rRNA copy numbers in soil X decreased, while those in soil Y did not.

**Figure 7.**
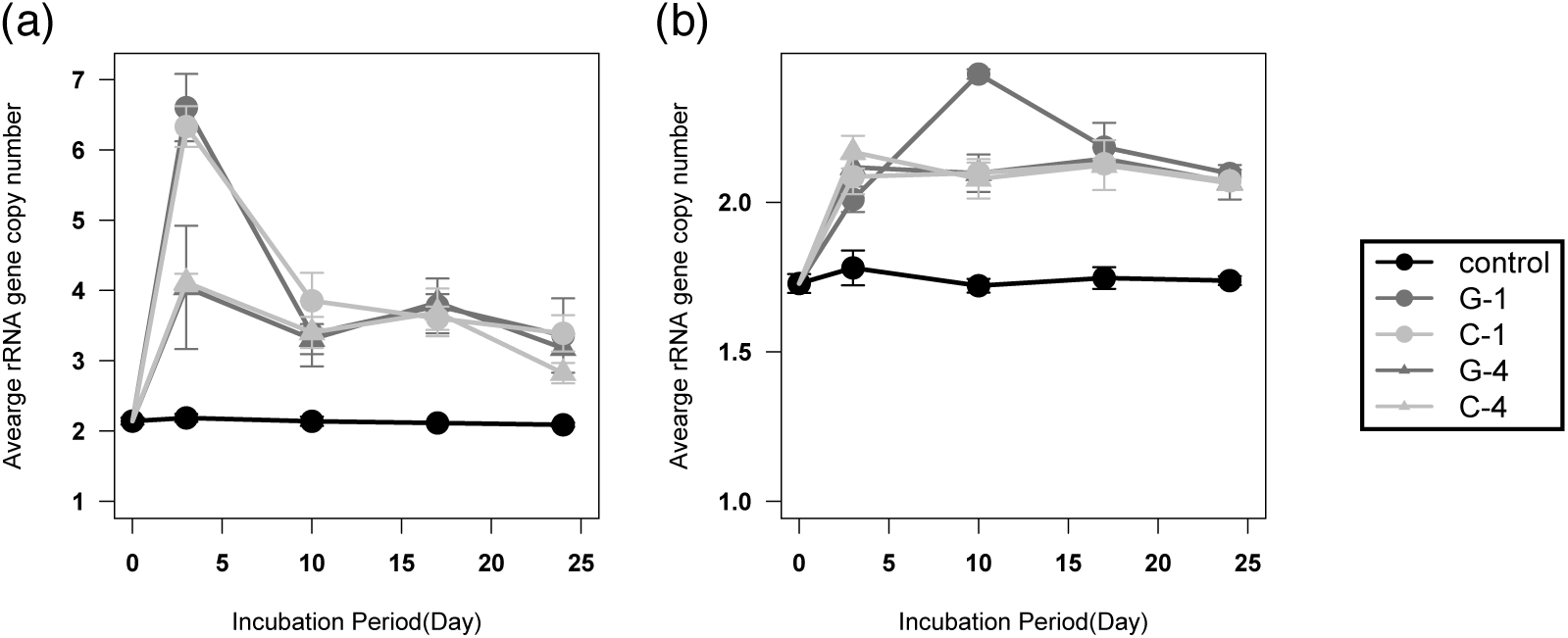
Transition of average rRNA gene copy number of observed OTUs in soil X (a) and soil Y (b), weighted by read counts. Presented are the average of three replicates.

## 4. Discussion

Overall, soil DOC concentration after 24 hours of incubation was overall higher in soil Y than soil X. This indicates that microbial carbon intake was slower in soil Y than in soil X, which means that soil Y was under a stark phosphorus limitation. This is in accordance with a lower concentration of available phosphorus in soil Y than soil X, and prevailing phosphorus-limiting conditions in forest soils in general (Kunito et al., 2012; Noyce et al., 2015).

Soil enzyme activity was also in accordance with the phosphorus-limited nature of soil Y. Although both soils presented a rise in microbial phosphatase activity after carbon and nitrogen amendment, soil Y was unique in that little difference in phosphatase activity was observed between two levels of CN amendment. In other words, the CN amount required to make soil Y phosphorus-limited was smaller than that of soil X. Additionally, regarding the groups G-1 and C-1, the ACP and ALP activities gradually decreased in soil X while they did not in soil Y from day 3 to 24, which also can too be attributed to the difference in soil phosphorus availability. In soil X, it can be assumed that most of the carbon and nitrogen amendments were quickly taken up by microbes (within 24 hours after amendment). Conversely, in soil Y, phosphorus scarcity apparently slowed down the carbon and nitrogen usage, and thus higher phosphatase activities were maintained. All the above inference regarding enzymatic activities is in line with the presupposition of the resource allocation model for extracellular enzymatic synthesis (Sinsabaugh et al., 1994), which claims that low phosphorus availability induces preferential production of phosphatase by microbes. Contrasting with previous studies that proved this model to be applicable to various soils in stable conditions (Fujita et al., 2017), our results indicate that short-term responses after trophic perturbation also can be explained by the concept of resource allocation.

However, the influence of the type of amended carbon source was not straightforward. According to the resource allocation model, microbial phosphatase production was predicted to be lower in the cellobiose-amended group than in the glucose-amended group, because microbes in cellobiose-amended soils allocate more resources to BG production than those in the glucose-amended soils. Given that the total amount of usable resources is the same in both glucose- and cellobiose-amended soils (or higher in glucose-amended soils), the phosphatase activity should be lower in cellobiose-amended soils. Some of our results — the higher BG activity in cellobiose-amended groups — are in line with the resource allocation model. Contrary to the inference above, the relationship between phosphatase activity and the amended carbon source was not consistent. One possible explanation for this observation is that microbes in the cellobiose-amended samples compromised the production of some other extracellular enzymes rather than extracellular phosphatase, but this cannot be directly determined from our data.

The transition in microbial community structures presented fundamental differences between the two soils, which can also be attributed to the differences in nutrient stoichiometry. Over the course of 24 days of incubation, microbial communities in soil X gradually recovered to the initial state while those in soil Y did not. Here we point out two aspects of microbial community assembly: microbial life-history strategy and competitions among microbes.

Microbial life history strategy is often explained by its maximum growth rate. Fast-growers favor copiotrophic environments, while slow-growers can survive in and dominate oligotrophic environments. The OTU annotated as the order *Bacillales*, which prominently contributed to community assembly in soil X, is a typical spore-forming fast-grower (Vos et al. eds., 2009). Moreover, recent studies suggest that rRNA gene copy number also represents microbial life history strategy (Roller et al., 2016). Given that microbes with a high copy number of rRNA genes are generally copiotrophic and present high growth rates (Klappenbach et al., 2000), soil X was presumably a more favorable environment to copiotrophic microbes than soil Y. To ensure rapid growth, phosphorus availability is required in addition to a high rRNA gene copy number, because phosphorus is a major component of nucleic acids and cell membranes. It can be assumed that the relatively high phosphorus availability in soil X drove the proliferation of fast-growers.

As suggested in a previous study (Delgado-Baquerizo et al., 2016), phosphorus limitation induces microbial competition and reduces community diversity. Thus, we assume that the continuous drop in community evenness in soil Y, compared with that of soil X, can be interpreted as the outcome of competition enhanced by stark phosphorus scarcity. Although our 16S rRNA gene sequencing results do not directly prove phosphorus competition because adaptation to phosphorus scarcity is not a phylogenetically conserved trait (Martiny et al., 2015), the inference above coincides with a prolonged and gradual increase in phosphatase activities in soil Y.

Collectively, our results show that the microbial community response to CN input was consistent with soil nutrient stoichiometry in both function and structure. Although soil X showed more drastic changes than soil Y in the very short term (up to three days), this effect did not last long and the community gradually returned to the initial state. By contrast, in soil Y, where phosphorus is highly depleted compared with soil X, CN amendment had a prolonged impact on the microbial community. These observations indicate that our initial hypothesis should be partially modified: strong phosphorus limitation results in prolonged, rather than drastic, changes in the microbial community.

## Data Availability

Nucleotide sequences obtained in this study will be made publicly available at the time of publication in a refereed journal.

## Acknowledgements

This study was supported by JSPS KAKENHI (Grant-in-Aid for Scientific Research (B)) No.26292035. The authors thank Kazuo Isobe (University of Tokyo) for advices in performing the overall research, and Hitoshi Moro (Shinshu University) for aid in soil sampling. The authors declare no conflict of interests.

## Author contributions

TK and SO conceived of the study. KM, RM and YM performed the experiments. KM, RM, TK, KS and SO interpreted the data. KM wrote the manuscript.

---

**Supplementary Figure 1.**
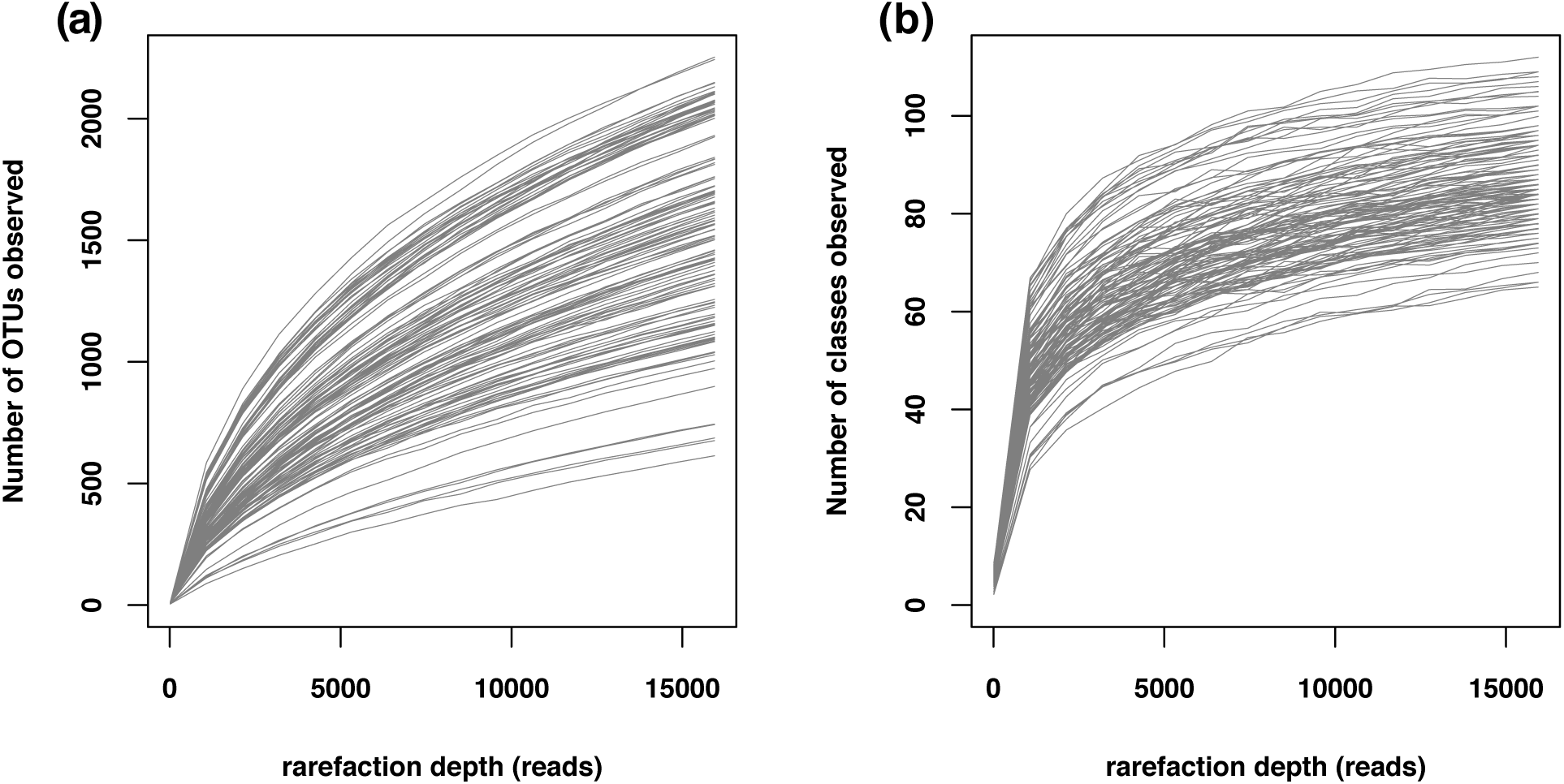
Rarefaction curves of 16S amplicons, indicating the number of observed (a) operational taxonomic units (OTUs) and (b) classes.

**Supplementary Figure 2.**
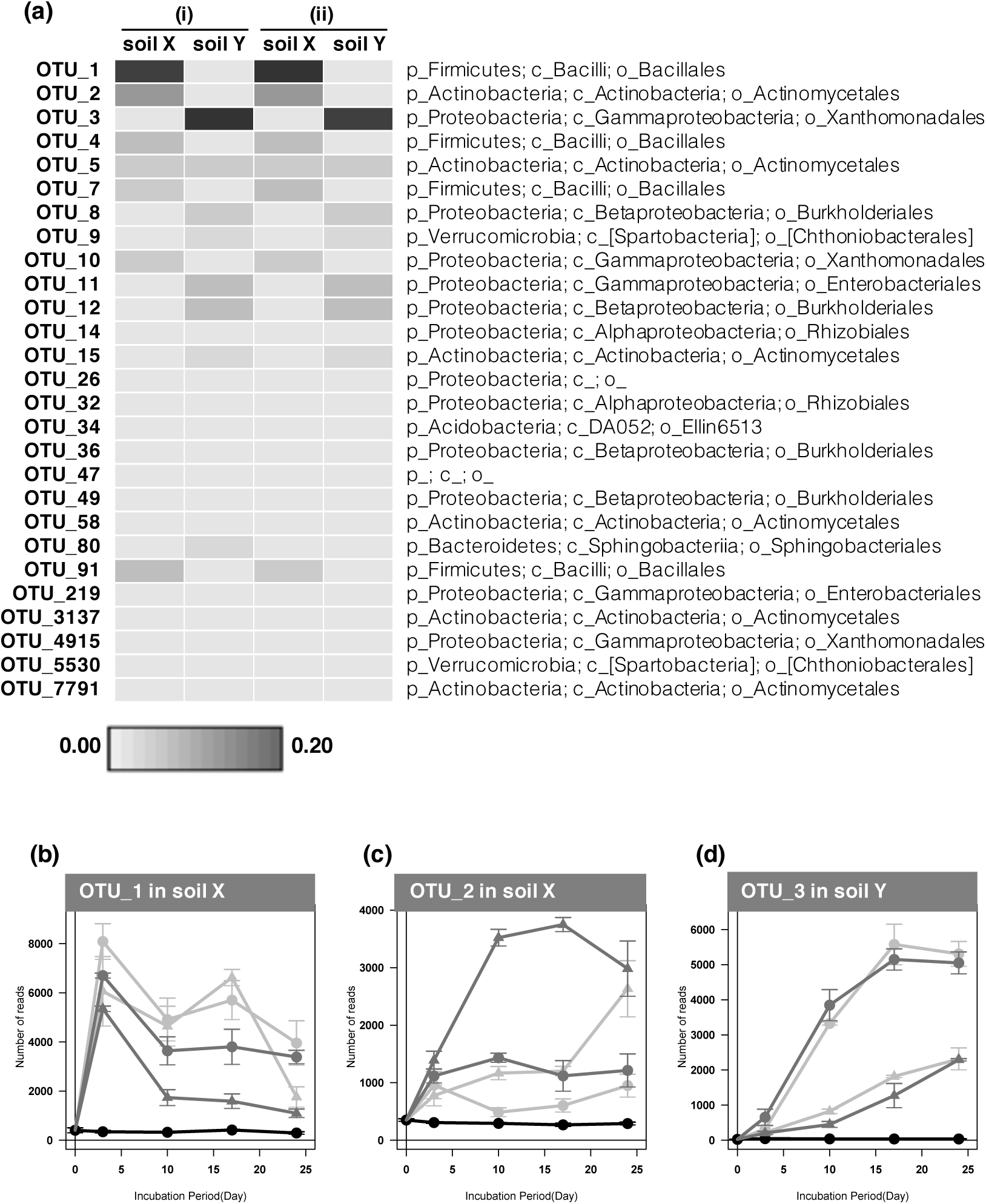
(a) The contribution of each OTU to the community structure dissimilarity between: (i) glucose-amended soils and cellobiose-amended soils (G-1+G-4 vs C-1+C-4) and (ii) CN amendment patterns (G-1+C-1 vs G-4+C-4), calculated based on SIMPER analysis. Twenty-seven OTUs, whose cumulative contribution rate is above 50% in at least one of the four analyses, are presented. (b-d) Relative abundance of three OTUs that highly contribute to overall community structure dissimilarity.

## References

Buscardo, E., Geml, J., Schmidt, S.K., Silva, A.L.C., Ramos, R.T.J., Barbosa, S.M.R., et al. (2018) Of mammals and bacteria in a rainforest: Temporal dynamics of soil bacteria in response to simulated N pulse from mammalian urine. Funct. Ecol. 32: 773–784.

Caporaso, J.G., Kuczynski, J., Stombaugh, J., Bittinger, K., Bushman, F.D., Costello, E.K., et al. (2010) QIIME allows analysis of high-throughput community sequencing data. Nat. Methods 7: 335–336.

Caporaso, J.G., Lauber, C.L., Walters, W.A., Berg-Lyons, D., Huntley, J., Fierer, N., et al. (2012) Ultra-high-throughput microbial community analysis on the Illumina HiSeq and MiSeq platforms. ISME J. 6: 1621–1624.

Caporaso, J.G., Bittinger, K., Bushman, F.D., Desantis, T.Z., Andersen, G.L., and Knight, R. (2010) PyNAST: A flexible tool for aligning sequences to a template alignment. Bioinformatics 26: 266–267.

Castle, S.C., Sullivan, B.W., Knelman, J., Hood, E., Nemergut, D.R., Schmidt, S.K., and Cleveland, C.C. (2017) Nutrient limitation of soil microbial activity during the earliest stages of ecosystem development. Oecologia 185: 513–524.

Delgado-Baquerizo, M., Reich, P.B., Khachane, A.N., Campbell, C.D., Thomas, N., Freitag, T.E., et al. (2017) It is elemental: soil nutrient stoichiometry drives bacterial diversity. Environ. Microbiol. 19: 1176–1188.

Edgar, R.C. (2013) UPARSE: highly accurate OTU sequences from microbial amplicon reads. Nat. Methods 10: 996–998.

Edgar, R.C. (2010) Search and clustering orders of magnitude faster than BLAST. Bioinformatics 26: 2460–2461.

Fierer, N. (2017) Embracing the unknown: Disentangling the complexities of the soil microbiome. Nat. Rev. Microbiol. 15: 579–590.

Fujita, K., Kunito, T., Moro, H., Toda, H., Otsuka, S., and Nagaoka, K. (2017) Microbial resource allocation for phosphatase synthesis reflects the availability of inorganic phosphorus across various soils. Biogeochemistry 136: 325–339.

Griffiths, B.S. and Philippot, L. (2013) Insights into the resistance and resilience of the soil microbial community. FEMS Microbiol. Rev. 37: 112–129.

Klappenbach, J.A., Dunbar, J.M., Thomas, M., and Schmidt, T.M. (2000) rRNA operon copy number reflects ecological strategies of bacteria. Appl. Envir. Microbiol. 66: 1328–1333.

Kunito, T., Tsunekawa, M., Yoshida, S., Park, H.-D., Toda, H., Nagaoka, K., and Saeki, K. (2012) Soil properties affecting phosphorus forms and phosphatase activities in Japanese forest soils. Soil Sci. 177: 39–46.

Langille, M.G.I., Zaneveld, J., Caporaso, J.G., McDonald, D., Knights, D., Reyes, J.A., et al. (2013) Predictive functional profiling of microbial communities using 16S rRNA marker gene sequences. Nat. Biotechnol. 31: 814–821.

Lozupone, C. and Knight, R. (2005) UniFrac: a New Phylogenetic Method for Comparing Microbial Communities UniFrac: a New Phylogenetic Method for Comparing Microbial Communities. Appl. Environ. Microbiol. 71: 8228–8235.

Martiny, J.B.H., Jones, S.E., Lennon, J.T., and Martiny, A.C. (2015) Microbiomes in light of traits: A phylogenetic perspective. Science. 350: aac9323.

McDonald, D., Price, M.N., Goodrich, J., Nawrocki, E.P., Desantis, T.Z., Probst, A., et al. (2012) An improved Greengenes taxonomy with explicit ranks for ecological and evolutionary analyses of bacteria and archaea. ISME J. 6: 610–618.

Murphy, J. and Riley, J.P. (1962) A modified single solution method for the determination of phosphate in natural waters. Anal. Chem. Acta 27: 31–36.

Nemergut, D.R., Knelman, J.E., Ferrenberg, S., Bilinski, T., Melbourne, B., Jiang, L., et al. (2016) Decreases in average bacterial community rRNA operon copy number during succession. ISME J. 10: 1147–1156.

Noyce, G.L., Basiliko, N., Fulthorpe, R., Sackett, T.E., and Thomas, S.C. (2015) Soil microbial responses over 2 years following biochar addition to a north temperate forest. Biol. Fertil. Soils 51: 649–659.

Pagaling, E., Strathdee, F., Spears, B.M., Cates, M.E., Allen, R.J., and Free, A. (2014) Community history affects the predictability of microbial ecosystem development. ISME J. 8: 19–30.

Price, M.N., Dehal, P.S., and Arkin, A.P. (2009) Fasttree: Computing large minimum evolution trees with profiles instead of a distance matrix. Mol. Biol. Evol. 26: 1641–1650.

R Core, T. (2017) R: A language and environment for statistical computing. R Foundation for Statistical Computing, Vienna, Austria. URL https://www.R-project.org/.

Roller, B.R.K., Stoddard, S.F., and Schmidt, T.M. (2016) Exploiting rRNA operon copy number to investigate bacterial reproductive strategies. Nat. Microbiol. 1: 1–7.

Saeki, K., Sakai, M., and Kunito, T. (2012) Effect of α-casein on DNA adsorption by Andosols and by soil components. Biol. Fertil. Soils 48: 469–474.

Sinsabaugh, R.L. and Moorhead, D.L. (1994) Resource allocation to extracellular enzyme production: A model for nitrogen and phosphorus control of litter decomposition. Soil Biol. Biochem. 26: 1305–1311.

Tabatabai, M.A. (1994) Soil enzymes. In, Weaver, R., Angle, S., Bottomley, P., Bezdicek, D., Smith, S., Tabatabai, M.A., and Wollum, A. (eds), Methods of Soil Analysis, Part 2, Microbiological and Biochemical Properties. Soil Science Society of America, Madison, Madison, Wisconsin, pp. 775–833.

Truog, E. (1930) The determination of the readily available phosphorus of soils. J. Am. Soc. Agron. 22: 874–882.

Vos, P., Garrity, G., Jones, D., Krieg N. R., Ludwig, G., Rainey, F. A., Shleifer, K-H., and Whitman W. B. (eds) Bergey’s Manual of Systematic Bacteriology. Volume Three: The Firmicutes. 2nd edn (Springer, 2009).

Wang, Q., Wang, S., He, T., Liu, L., and Wu, J. (2014) Response of organic carbon mineralization and microbial community to leaf litter and nutrient additions in subtropical forest soils. Soil Biol. Biochem. 71: 13–20.

Wang, Q., Garrity, G.M., Tiedje, J.M., and Cole, J.R. (2007) Naive Bayesian classifier for rapid assignment of rRNA sequences into the new bacterial taxonomy. Appl. Environ. Microbiol. 73: 5261–5267.

